# Assessing Habitat Dynamics and Land-Use Patterns in the Amazon Forest Using Satellite Imagery

**DOI:** 10.64898/2026.02.17.706446

**Authors:** Elizabeth Rentería

## Abstract

Tropical forests, particularly the Amazon, play a critical role in global ecosystems by providing essential services such as climate regulation, carbon sequestration, and biodiversity conservation. However, these ecosystems are increasingly threatened by deforestation and land-use changes driven by agriculture, livestock farming, and other anthropogenic activities. This study investigates habitat composition and temporal changes in Tailândia (Pará-Brazil), using high-resolution satellite imagery. Data from 2013 to 2023 were analyzed across 18 research plots and a broader expanded zone to identify patterns of land-use transformation. Results reveal the dominance of Forest Formation habitats, alongside significant increases in Pastures and Oil Palm Crops. Clustering analysis highlighted ecological heterogeneity, with intact forests and heavily altered plots demonstrating varied conservation needs. Results also forecast a 13% decline in forest cover and a 32% rise in pasture areas over the next five years. The findings underscore the urgent need for targeted conservation strategies, robust environmental policies, and sustainable land-use practices. This research demonstrates the utility of remote sensing for large-scale ecological monitoring and its potential to inform effective conservation efforts.

## Introduction

Tropical forests are invaluable to global ecosystems, providing essential services that reach beyond their geographic boundaries. Among the most significant of these services is climate change mitigation, as tropical forests store and sequester substantial amounts of carbon, helping to stabilize and reduce atmospheric CO_2_ levels (Houghton et al., 2015). They also contribute to maintaining rainfall patterns, reduce drought risk, and support stable water levels by retaining moisture, providing clean water for human consumption (Watson et al., 2018; Chandrathilake, 2022). Healthy forests also help control outbreaks of vector-borne diseases. Additionally, tropical forests are home to numerous indigenous communities who strongly depend on their resources (Myers et al., 2009; Watson et al., 2018). These forests are also biodiversity hotspots, they shelter approximately two-thirds of the world’s species, making them the most biodiverse ecosystems at every taxonomic level. This biodiversity underscores the need for focused conservation efforts, as these ecosystems are critical for sustaining global species diversity (Merritt et al., 2019).

Along with land conversion for cattle ranching and logging, agricultural expansion is a leading driver of biodiversity loss, disrupting the ecological functions that many organisms provide (Nunes et al., 2022; Ewers, 2015). Primary forests are especially vulnerable, as they are increasingly cleared to meet rising global demand for agricultural land. Tropical areas are projected to become major spots for agriculture, as other regions, decrease their land use for this purpose (Gibbs et al., 2010). This damage not only impacts the targeted areas but also neighbouring forests through edge effects and fragmentation (Lewis et al., 2015). Forest fragmentation and edge further exacerbates these impacts by causing the local extinction of habitat-specialist species, and even more adaptable species are at risk as they encounter thresholds of habitat size and quality that they cannot survive (Hanski, 2015). Preserving primary forests is vital because secondary forests, while beneficial, cannot fully replicate the biodiversity and ecosystem services of primary forests (Gibson, 2011). Understanding habitat dynamics at both local and regional scales is crucial for effective conservation strategies.

This study seeks to corroborate the effectiveness of remote sensing in monitoring habitat dynamics and land-use changes in the Amazon rainforest. Remote sensing offers consistent and scalable methods to study vast areas. The Amazon’s challenging terrain makes traditional field-based surveys impractical; remote sensing addresses this by providing frequent, consistent observations, enabling multi-scale analysis of dynamic changes (Almeida et al., 2016; Souza et al., 2013; Souza et al., 2023). This study investigates habitat composition and temporal changes in Tailândia (Pará-Brazil), using high-resolution satellite imagery and advanced analytical techniques. By integrating temporal and spatial datasets, this study highlights how these tools reveal ecological patterns and inform conservation practices.

In this study, satellite imagery provided detailed classifications of 11 habitat types across 11 years. The area was divided in 18 research plots with a 2 km radius each and a broader expanded zone with a 30 km radius. This multi-scale approach enabled the identification of both localized patterns, such as habitat diversity within individual plots, and larger-scale trends, including the dominance of forest formations and the progressive expansion of Oil palm crops and pastures. Such insights would be difficult to achieve through ground-based methods alone, underscoring the importance of satellite data as a cornerstone of ecological research. The temporal analysis over a decade (2013-2023) provided insights into long-term trends, underscoring remote sensing’s utility in addressing complex environmental questions. This study aims to document clear land-use changes in the broader study area while detecting subtler shifts within individual plots, laying the groundwork for future ecological research in this region.

## Methods

### Data Collection and description

Data was collected using Google Earth Engine (GEE), specifically the Mapbiomas User Toolkit (Collection 9: 1985 through 2023; Landsat 5, 7, 8, and 9), to generate maps of Brazil. This provided access to high-resolution satellite imagery and environmental datasets spanning 11 years.

The study area is Tailândia, Pará, Brazil (2°30’23.8”S 48°45’57.0”W), comprising 18 distinct research plots within the Amazon forest. Each plot has a radius of 2 km and includes a unique combination of habitat types. There are 11 habitat types, with each plot containing between 1 and 6 types. Additionally, an expanded zone was established, encompassing all 18 plots and the surrounding land. This expanded zone, with a 30 km radius, offers a broader context for understanding land use patterns and habitat composition in the region (Figure 1). The expanded zone includes 13 habitat types. Analysing this dataset helps identify larger-scale land use changes and patterns that may not be apparent from individual plots. Satellite images from 2013 to 2023 were used to extract measurements for each habitat type.

**Fig 1.**
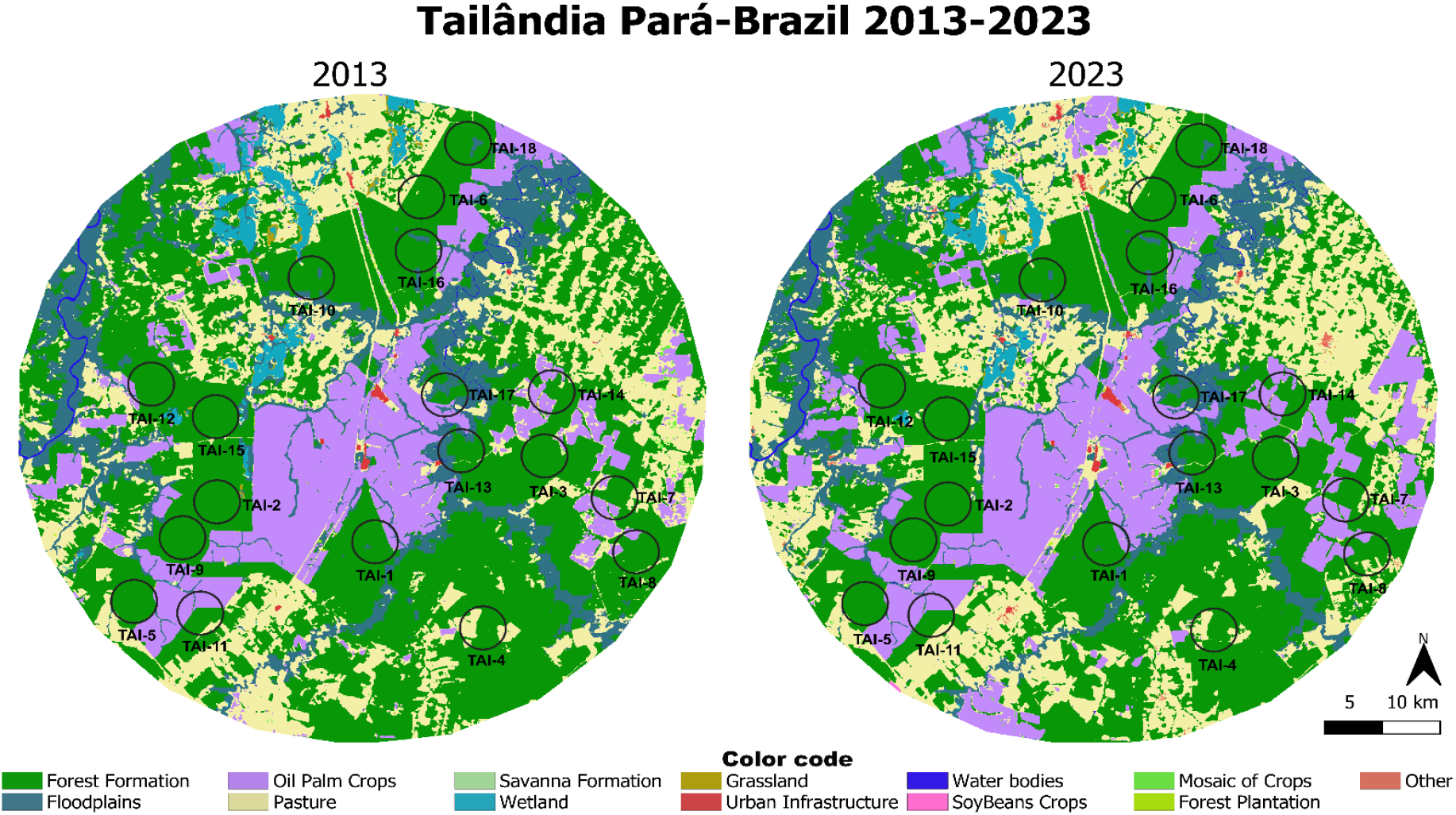
Map of the study area (Tailândia, Pará-Brazil 2013-2023), showing the location of the 18 plots and the expanded zone. Zones determined with Mapbiomas User Toolkit-Collection 9. Image created with QGIS.

The habitat types can be divided into natural and anthropogenic habitats. Forest Formation, Floodplains, Grasslands, Savannas, Wetlands are natura habitats. While the others are from anthropogenic origin; except from water bodies, that can be both.

### Data Analysis

Data analysis was conducted using Python (version 3.13.1). Initially, K-means clustering, an unsupervised machine learning algorithm, was applied to group research plots based on their habitat composition similarity. This approach identified spatial patterns and areas of interest with similar ecological characteristics. While clustering provided initial insights, it lacked detailed clarity. Consequently, heatmaps were created to visualize habitat distribution more effectively. Heatmaps illustrate spatial variations in habitat composition, enabling a detailed landscape analysis.

To determine the statistical significance of observed differences in the heatmaps, an Analysis of Variance (ANOVA) was conducted. This test assessed whether variations in habitat composition across different plots were significant or due to random variation. ANOVA was paired with a Tukey test to identify specific variables driving these variations, validating the robustness of the results and understanding regional environmental variability.

To explore potential future trends in habitat dynamics, a time series analysis using the ARIMA (AutoRegressive Integrated Moving Average) model was performed. This model forecasts future habitat composition changes based on historical satellite data, capturing patterns and trends over time. A five-year forecast was generated. Additionally, linear regression was applied to the expanded zone to examine relationships between the areas of the four most dominant habitat types and their changes over the past 11 years. This regression quantified trends in land use and habitat composition, highlighting potential ecological change drivers such as deforestation or urbanization.

## Results

### Cluster Data exploration

K-means clustering revealed distinct ecological groupings among the 18 plots (Fig. 2). Plots dominated by “Forest Formation” formed separate clusters, emphasizing the predominance of natural forest ecosystems in these areas, such as plot TAI_02 and TAI_09, that only have a single habitat type (“Forest Formation”). In contrast, plots with a diverse number of habitats clustered differently, like plot TAI_10 and TAI_13, which each have six different habitat types. This variation in clustering indicates differing habitat type diversity among plots. Clustering of data from the expanded zone showed a clear dominance of the “Forest Formation” habitat type, followed by “Pastures” and “Oil Palm Crops” (Fig.2).

**Fig 2.**
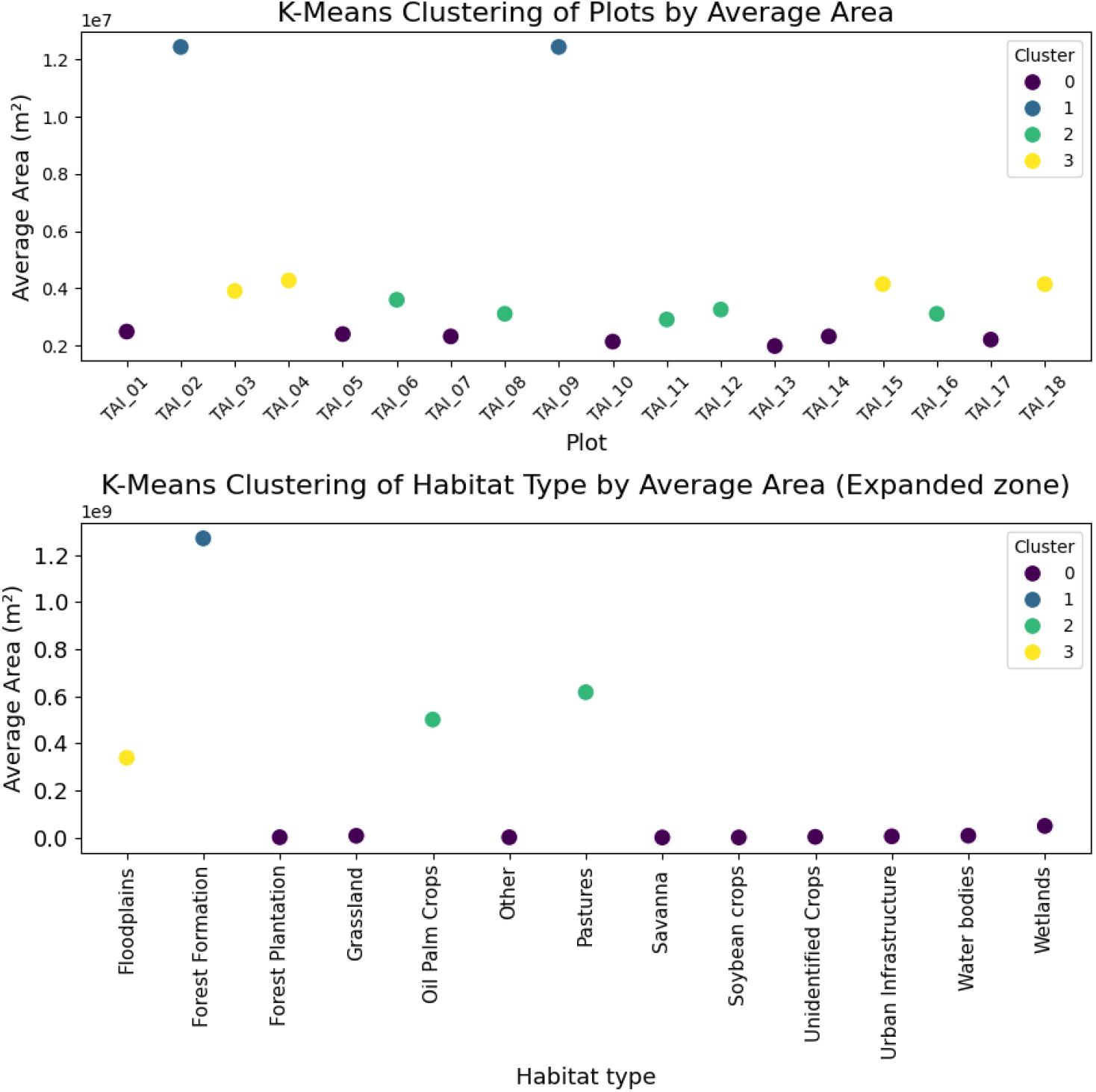
K-means clustering. The upper plot represents the levels of area homogeneity or heterogeneity of the each of the 18 plots. Cluster 1 is the more homogenous, so they have more area because it’s only distributed in 1 habitat type. Cluster 0 are plots that have their area divided into five or six habitat types. Cluster 2, three-four habitat types. Cluster 3, two-three habitat types. The lower plot indicates which habitat type has a higher area in the expanded zone.

### Habitat type area distribution (Heatmaps)

Heatmaps provided detailed visualizations of habitat composition in each plot and annually (Fig. 3). “Forest Formation” dominated most plots, often comprising over 70% of the area. “Oil Palm Crops” consistently appeared as a secondary habitat type in specific plots, contributing up to 25% in some areas. Most habitats contributed with less than 1% to the plots. The expanded zone’s heatmap focused on habitat type size distribution over the last 11 years. The expanded zone’s heatmap focused on habitat type size distribution over the last 11 years. Temporal heatmaps revealed a higher proportion of “Forest Formation” across years, with a small but consistent decline. “Oil Palm Crops” and “Pastures” showed a clear upward trend, indicating progressive forest clearance over the decade. “Floodplains” also significantly contributed to the area’s composition. Other habitats remained constant with minimal contribution.

**Fig 3.**
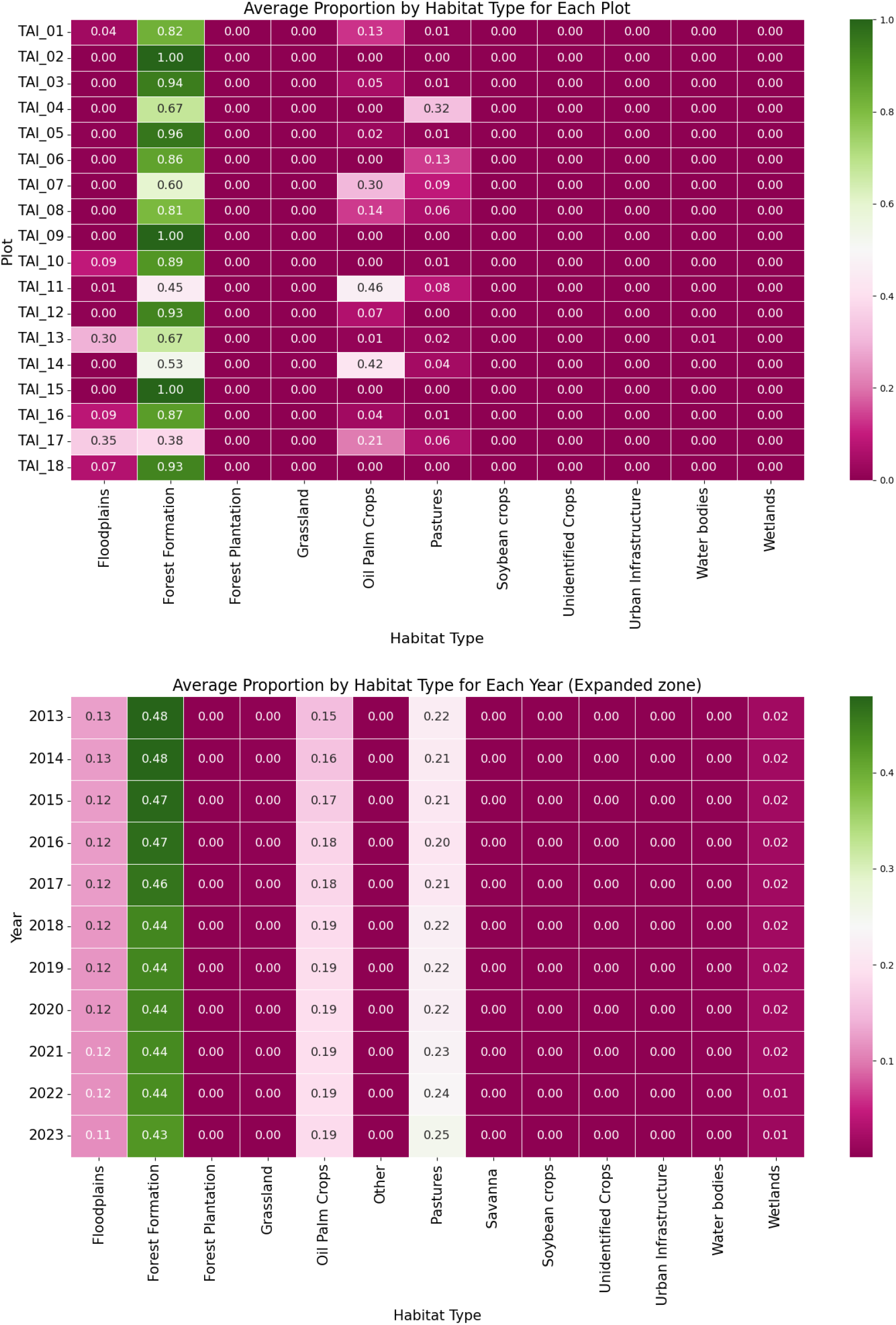
Heatmaps to visualize the distribution percentage of habitat types. The upper plot shows the distribution of habitats in total in the 11 years. The lower plot shows the distribution percentage of habitats of the expanded zone, showed annually.

### ANOVA and Tukey test

ANOVA results confirmed significant variations in habitat composition across the 18 plots (F-statistic: 464.76, p-value: 1.64 × 10^-321^). Tukey’s test helped identify “Forest Formation”, as the only habitat type significantly higher than the other habitats (Fig. 4). In the expanded zone, ANOVA showed significant variations in habitat composition (F-statistic: 3518.15, p-value: 5.48 × 10^-157^). In this case, the habitats that influence the variation were “Forest Formation”, “Oil Palm Crops”, “Pastures” and “Floodplains” (Fig. 4).

**Fig 4.**
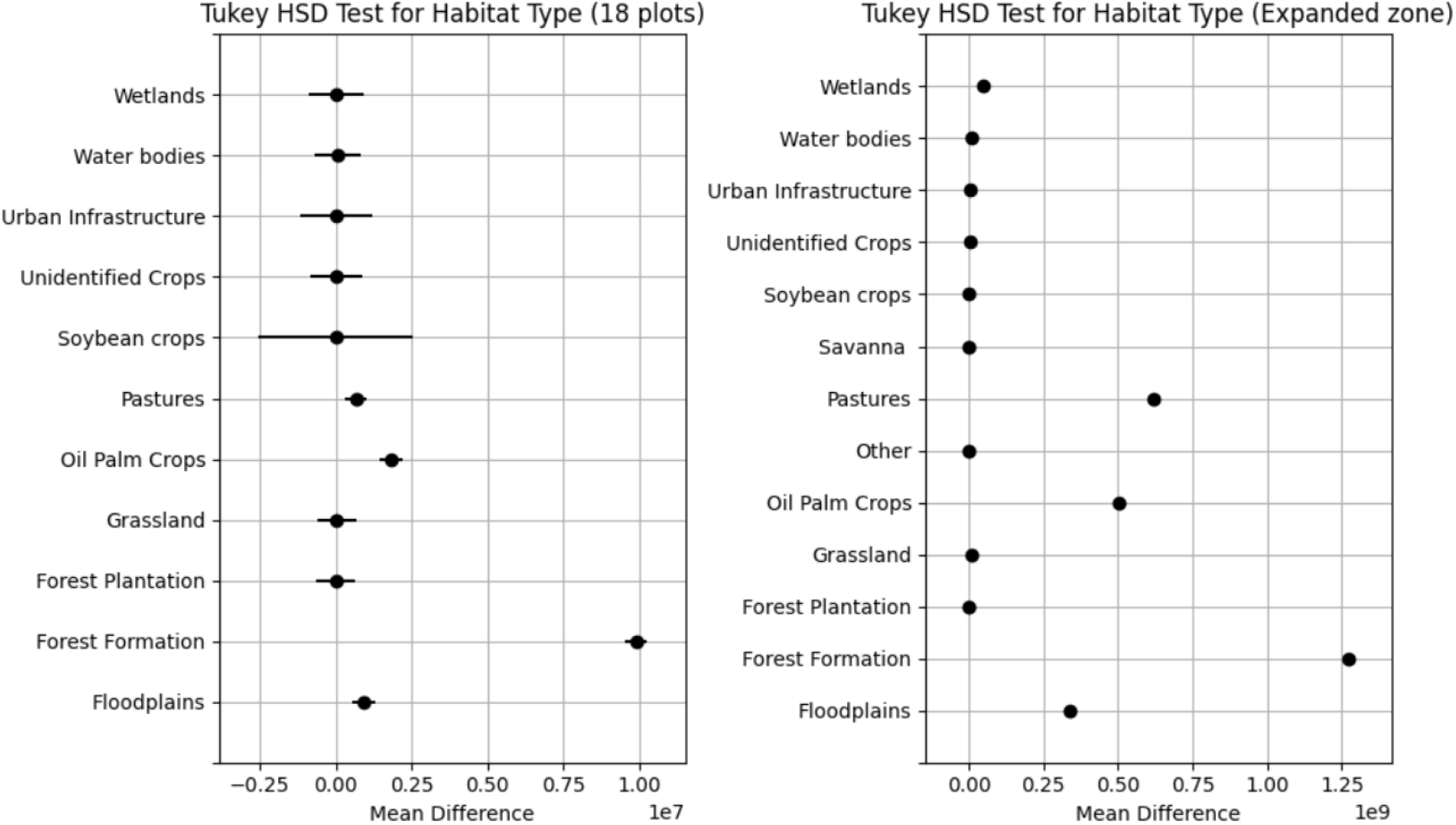
Tukey test plots. Left plot shows the difference of habitat types in the 18 plots. The right plot shows the difference in the expanded zone.

### Time Analysis and 5-year forecast of the expanded zone

The time analysis and forecast were only performed on the four largest habitats (Fig. 5). The forecast showed that the area of the habitat “Forest Formation” expected to decrease around 13% of its 2023 extent (1.193 x10^9^ m2) in the next 5 years. The habitat “Pastures” is expected to have a grow of about 32 % of its 2023 extent (7.037x10^8^ m2) in the next 5 years. The habitat “Oil Palm Crops” is expected to have a decrease of about 11% of its 2023 extent (5.219x10^8^ m2) in the next 5 years. The habitat “Floodplains” is expected to have a decrease of about 9% of its 2023 extent (3.191x10^8^ m2) in the next 5 years. The forecasts accuracy was ensured by fitting an ARIMA model to training data, providing reliable predictions (Fig. 6).

**Fig 5.**
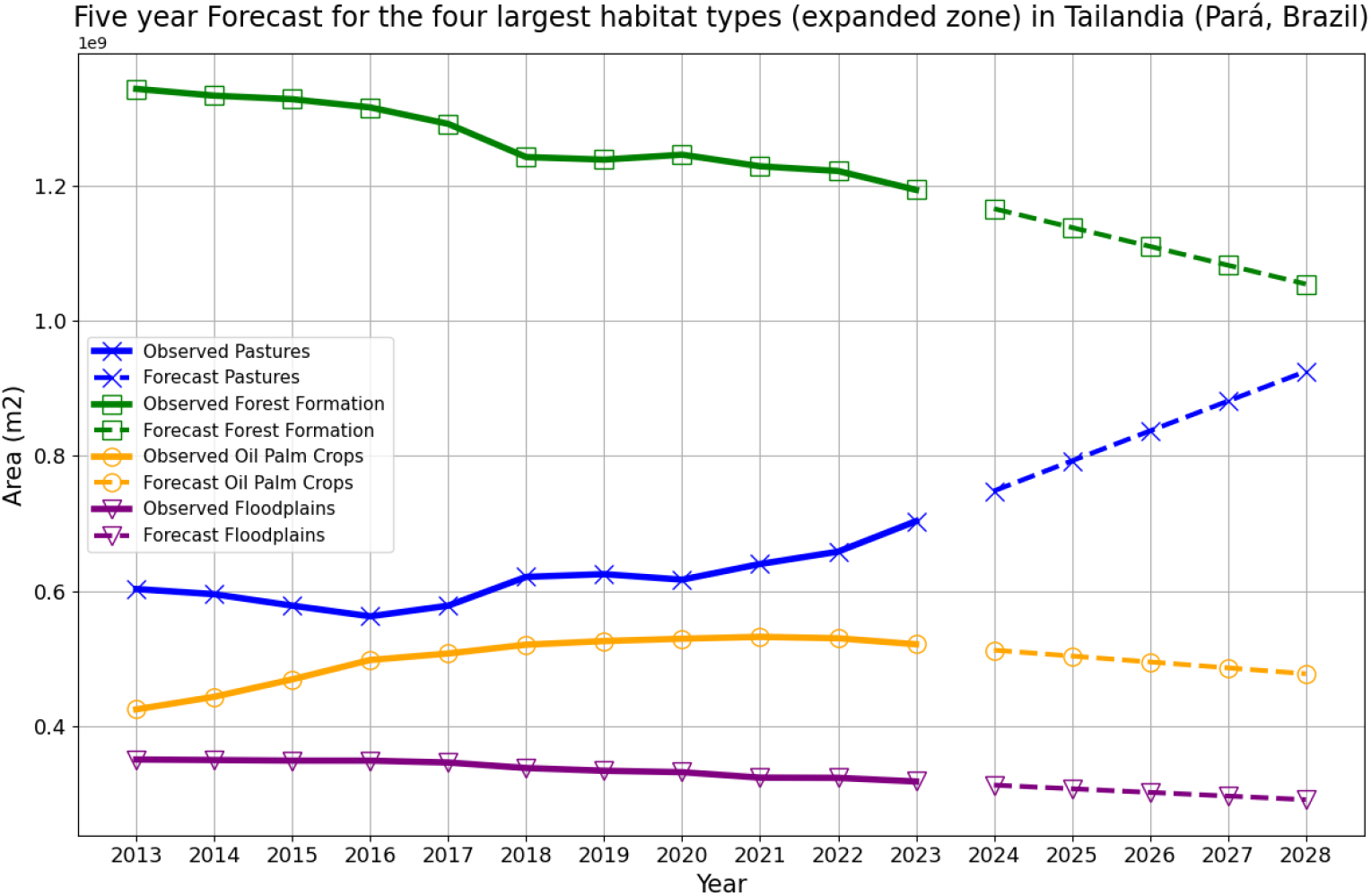
Five-year forecast of the four largest habitat types in the expanded zone.

**Fig 6.**
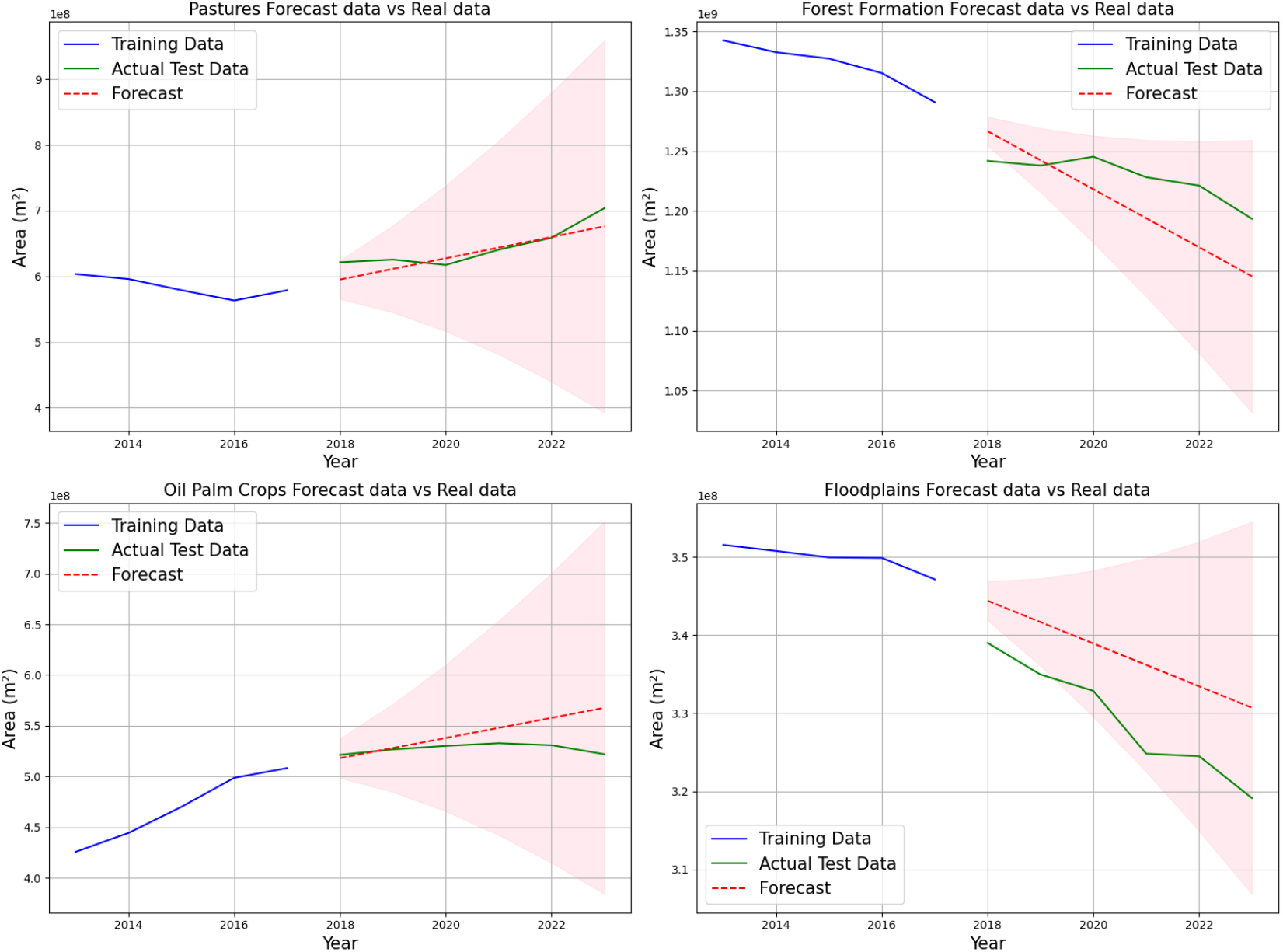
Fitted ARIMA model on training data (50% of data), to test accuracy of forecast vs actual test data (50% of data) of “Pastures”, “Forest Formation”, “Oil Palm Crops” and “Floodplains”.

### Linear regression

The OLS regression was applied to the four largest habitats (Fig. 7) in the expanded zone. The results for “Forest Formation” showed a significant negative trend, p-value: 0.000; R-squared: 0.939; coefficient: -1.528x10^7^; Prob (F-statistic): 9.02x10^-7^; diagnostic tests generally suggest the residuals behave well, with no major deviations from normality or strong autocorrelation (Omnibus test; Durbin Watson; Jarque-Bera). “Oil Palm Crops” demonstrated a positive trend p-value 0.000; R-squared: 0.772; coefficient: 9.969x10^6^; Prob (F-statistic): 0.0003; diagnostic tests generally good, but the Durbin-Watson statistic suggests some concern with autocorrelation, which could affect the model’s validity. Similarly, “Pastures” displayed a significant positive trend, p-value: 0.002; R-squared: 0.664; coefficient: 9.929x10^6^; Prob (F-statistic): 0.002; diagnostic tests generally good, but the Durbin-Watson statistic suggests a small concern with autocorrelation. “Floodplains” also showed a significant negative trend, p-value: 0.000; R-squared: 0.942; coefficient:-3.537x10^6^; Prob (F-statistic): 7.30x10^-7^; diagnostic tests generally suggest the residuals behave well, with no deviations from normality or strong correlation. All the regression were significative, although for the ones with a lower R-squared and doubts of corelation there may be other factors or variables that could improve the model’s explanatory power.

**Figure 7.**
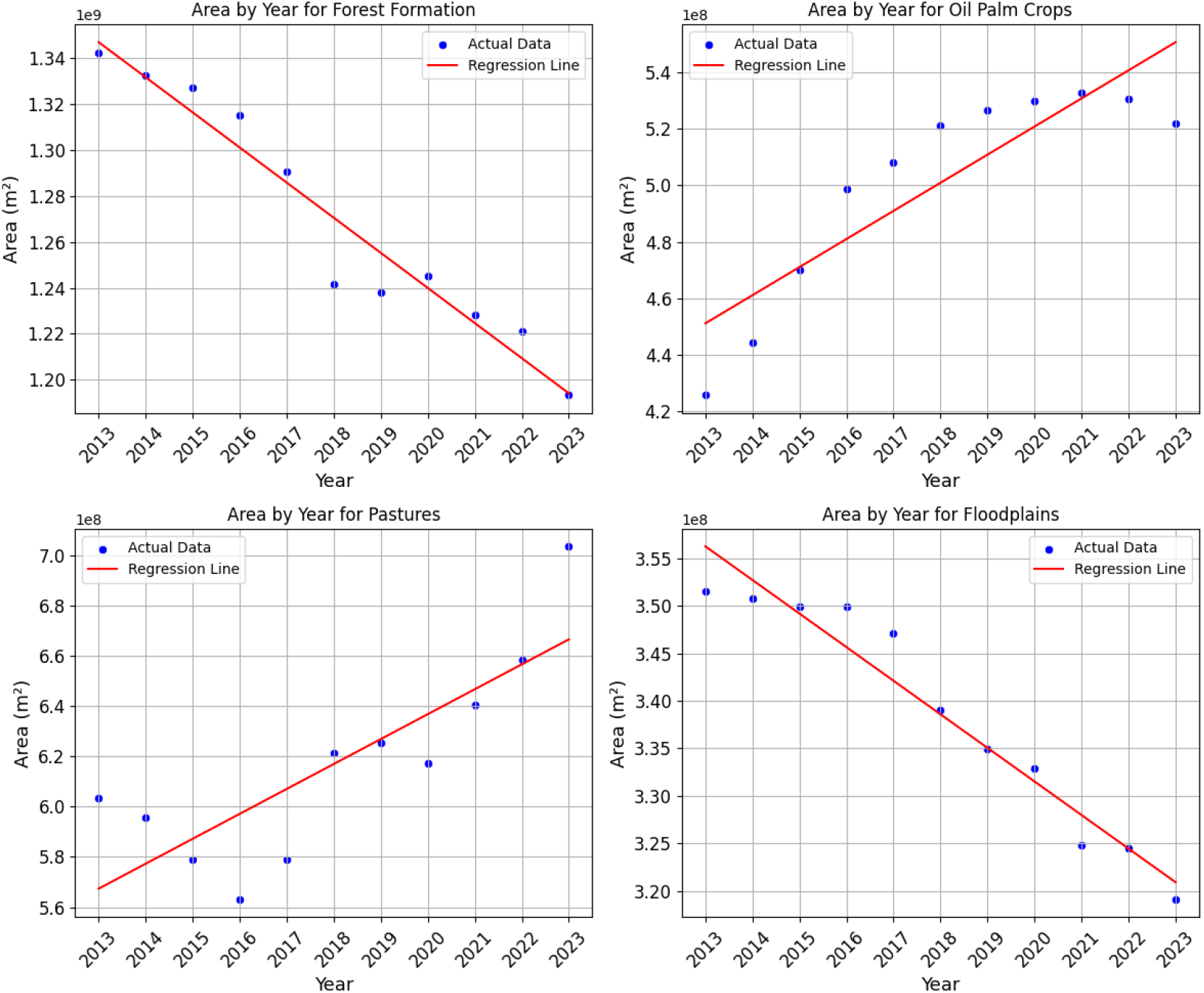
Linear regression graphs of the four largest habitat types in the expanded zone comparing their area across eleven years.

## Discussion

This study highlights the effectiveness of remote sensing in analyzing habitat dynamics and land-use changes in the Amazon rainforest. Remote sensing’s ability to consistently capture high-resolution spatial and temporal data makes it indispensable for studying vast ecosystems like the Amazon, where ground-based methods can be impractical or not possible (Luque et al., 2018). The results of this study provide interesting insights into the dynamic interplay between natural ecosystems and anthropogenic activities in the Amazon rainforest.

The predominance of natural occurring forest across the studied plots and the expanded zone underscores the Amazon’s role as a globally significant carbon sink and biodiversity hotspot (Antonelli et al., 2018; Gatti et al., 2021; Nogueira et al., 2014). However, the gradual but consistent expansion of Oil Palm Crops and Pastures reveals escalating pressures from agricultural and economic activities, with agricultural expansion being a major driver of deforestation in the Amazon (Pendrill et al., 2022). These findings align with previous studies that highlight the Amazon as a contested landscape, where economic drivers often conflict with ecological preservation (Carvalho et al., 2019; Silva et al., 2021).

The clustering analysis showed patterns of ecological heterogeneity among the plots. For instance, plots such as TAI_02 and TAI_09, dominated by a single habitat type, reflect relatively intact forest ecosystems, serving an important control area for comparing ecological data. In contrast, plots like TAI_10 and TAI_13, characterized by a mix of up to six habitat types, highlight regions undergoing significant land-use transformations.

The temporal analysis revealed alarming trends in the Amazon’s forest cover. A projected 13% decline in “Forest Formation” over the next five years signals an accelerating rate of deforestation. This is particularly concerning given the Amazon’s critical role in regulating global climate patterns, storing carbon, and serving as a refuge for a substantial portion of the world’s biodiversity (Antonelli et al., 2018; Tonetti et al., 2022). Disrupting the Amazon’s ecological balance could result in changes that would take centuries to restore (Liebsch et al., 2008). Furthermore, most carbon stock is concentrated in primary forests (Mackey et al., 2020). The scale of deforestation has pushed the Amazon to the brink of losing its resilience, possibly already crossing it’s tipping point. The consequences of its degradation would be felt not just locally but globally (Boulton et al., 2022).

At the same time, the anticipated 32% growth in Pastures emphasizes the expansion of livestock farming and agriculture, key drivers of deforestation in Brazil. This shift raises concerns about soil degradation, greenhouse gas emissions, and the sustainability of such land-use practices. Pastures dominate over 60% of the deforested areas in the Amazon, with cattle ranching as the primary use (Carvalho et al., 2020; Espindola et al., 2012). Pará, in particular, has seen significant intensification of pastureland. Despite the Amazon being less profitable for ranching compared to other regions, it remains a focal point for cattle production (Bowman et al., 2012). Notably, a substantial portion of cattle ranching occurs illegally. The Brazilian Amazon includes roughly 500,000 km^2^ of undesignated public forests, over 5% of which were illegally deforested by 2018 without repercussions, leaving the remaining areas highly vulnerable (Azevedo-Ramos & Moutinho, 2018; zu Ermgassen et al., 2024).

On a more positive note, the stability and slow decline in oil palm production suggest a reduction in land exploitation for this crop. Tailândia, located within Pará, is a prominent oil palm production zone due to its ideal climatic conditions (Almeida et al., 2019). Over the past six years, oil palm cultivation has remained relatively stable at 19% of the total agricultural area, showing a slight declining trend. This stability may result from stricter regulations and initiatives like the Roundtable on Sustainable Palm Oil (RSPO), the enforcement of Brazil’s Forest Code, and biome-wide commitments to zero deforestation (Brandão et al., 2021; Garrett et al., 2019; Soterroni et al., 2018). However, it is crucial to prevent large-scale expansion of oil palm plantations, as they are linked to environmental degradation, loss of biodiversity, and negative impacts on air, water, and soil quality (Manhães et al., 2024).

Floodplains are also projected to experience a gradual but consistent decline. These forests are among the most endangered habitats in the Amazon. In some areas, as much as half of the floodplain forests have been cleared, primarily for agriculture and cattle ranching (Renó et al., 2011). These ecosystems are vital for maintaining the Amazon’s resilience. Their degradation is associated with increased wildfires and significant carbon release, particularly from peatlands (Flores et al., 2017). Disturbances in these areas can trigger large-scale ecological consequences that extend far beyond the Amazon (Correa et al., 2022).

Historically, deforestation in the Brazilian Amazon peaked in 2004, with a loss of 28,000 km^2^. This was followed by an 84% decline by 2012, when it hit an all-time low, thanks to the implementation of anti-deforestation policies by the then-government (Ritchie, 2023). This was thanks to anti deforestation initiatives implemented by the then government. When the political party change (Bolsonaro), so did the deforestation levels, in recent years it has double since 2012 (Pereira et al, 2019; Ritchie, 2023). This highlights the significant influence of government policies anti deforestation initiatives, which, continuity or discontinuity, can be highly dependent on political leadership (Silva et al., 2023). With the return of Lula’s administration in 2023, there is renewed hope for reversing these trends, as the government has pledged to end deforestation by 2030 (Ritchie, 2023).

The findings of this study extend beyond the Amazon, offering valuable lessons for other regions facing similar challenges. The observed patterns of habitat change underscore the universal tension between economic development and environmental conservation. By adopting the methodologies and insights presented here, other tropical regions can enhance monitoring efforts, mitigate habitat loss, and promote sustainable development strategies.

While this study provides a comprehensive overview of land-use changes over the past 11 years, the full ecological consequences of these changes require field-based research. Future studies should incorporate on-the-ground biodiversity assessments, soil health evaluations, and carbon stock analyses to fully understand the environmental impact of land conversion. As emphasized at the beginning of this study, these findings serve as baseline data for future ecological research in this region.

## Conclusions

This study highlights the critical importance of remote sensing for monitoring and understanding habitat dynamics and land-use changes in the Amazon rainforest. The findings reveal significant ecological trends: the dominance of “Forest Formation” habitats, the expansion of “Pastures,” and the stabilization of “Oil Palm Crops.” The projected 13% decline in forest cover over the next five years, coupled with a 32% increase in pasture areas, underscores the urgent need for conservation interventions. Deforestation trends threaten the Amazon’s role as a global carbon sink and biodiversity hotspot, with cascading effects on climate regulation, species survival, and ecosystem services. The stabilization of “Oil Palm Crops” suggests progress in regulatory efforts, but continued vigilance is needed to prevent further habitat degradation. Additionally, the decline of floodplains highlights the need to protect these critical ecosystems, which play an essential role in hydrological cycles and carbon storage.

## Recommendations

Incorporating additional data layers, such as socioeconomic indicators, climate models, and species distribution data, to capture the multifaceted drivers of habitat change. Expanding the temporal scope of analysis to include historical data prior to 2013, providing a longer-term perspective on land-use trends, and keep updating our data yearly. Additionally, while satellite imagery offers unparalleled spatial and temporal coverage, fieldwork efforts are essential to validate the accuracy of habitat classification and ensure the reliability of the results.

## References

Almeida A, Guimarães I, & Ferraz, S. 2019. Long-term assessment of oil palm expansion and landscape change in the eastern Brazilian Amazon. Land Use Policy, 90. 104321. 10.1016/j.landusepol.2019.104321

Almeida C, Coutinho A, Dalla J, Esquerdo M, Adami M, Venturieri A, Diniz C, Dessay N, Durieux L, & Gomes A. 2016. High spatial resolution land use and land cover mapping of the Brazilian Legal Amazon in 2008 using Landsat-5/TM and MODIS data. Acta Amazonica, 291: 291–302. 10.1590/1809-4392201505504

Antonelli A, Zizka A, Carvalho F, Scharn R, Bacon C, Silvestro D, & Condamine F. 2018. Amazonia is the primary source of Neotropical biodiversity. PNAS, 115(23): 6034–6039. 10.1073/pnas.1713819115

Azevedo-Ramos C & Moutinho P. 2018. No man’s land in the Brazilian Amazon: Could undesignated public forests slow Amazon deforestation? Land Use Policy, 73: 125–127. 10.1016/j.landusepol.2018.01.005

Boulton C, Lenton T, & Boers N. 2022. Pronounced loss of Amazon rainforest resilience since the early 2000s. Nature Climate Change, 12: 271–278. 10.1038/s41558-022-01287-8

Bowman M, Soares-Filho B, Merry F, Nepstad D, Rodrigues H & Almeida O. 2012. Persistence of cattle ranching in the Brazilian Amazon: A spatial analysis of the rationale for beef production. Land use policy, 29: 558–568

Carvalho W, Mustin K, Hilário R, Vasconcelos I, Eilers V & Fearnside P. 2019. Deforestation control in the Brazilian Amazon: A conservation struggle being lost as agreements and regulations are subverted and bypassed. Perspectives in Ecology and Conservation, 17. 10.1016/j.pecon.2019.06.002.

Carvalho R, de Aguiar A & Amaral S. 2020. Diversity of cattle raising systems and its effects over forest regrowth in a core region of cattle production in the Brazilian Amazon. Regional Environmental Change, 20: 44. 10.1007/s10113-020-01626-5

Chandrathilake T. 2022. The need of ecohydrological research in tropical forests for healthy watersheds. Journal of Tropical Forestry and Environment, 12(02). 10.31357/jtfe.v12i02.6346

Correa S, van der Sleen P, Siddiqui S, Bogotá-Gregory J, Arantes C, Barnett A, Couto T, Goulding M, & Anderson E. 2022. Biotic Indicators for Ecological State Change in Amazonian Floodplains. BioScience, 72(8): 753–768. 10.1093/biosci/biac038

Espindola G, de Aguiar A, Pebesma E, Câmara G, Fonseca L. 2012. Agricultural land use dynamics in the Brazilian Amazon based on remote sensing and census data. Applied Geography, 32: 240–252

Ewers R, Boyle M, Gleave R, Plowman N, Benedick S, Bernard H, Bishop T, Yahya B, Chey V, Chung A, Davies R, Edwards D, Eggleton P, Fayle T, Hardwick S, Rahman H, Kitching R, Khoo M, Luke S & Turner E. 2015. Logging cuts the functional importance of invertebrates in tropical rainforest. Nature Communications, 6: 6836. 10.1038/ncomms7836.

Flores B, Holmgren M, Xu C, van Nes E, Jakovac C, Mesquita R, & Scheffer M. 2017. Floodplains as an Achilles’ heel of Amazonian forest resilience. PNAS, 114(17): 4442–4446. 10.1073/pnas.1617988114

Gatti L, Basso L, Miller J, et al. 2021. Amazonia as a carbon source linked to deforestation and climate change. Nature, 595: 388–393. 10.1038/s41586-021-03629-6

Gibbs H, Ruesch A, Achard F, Clayton M, Holmgren P, Ramankutty N, & Foley J. 2010. Tropical forests were the primary sources of new agricultural land in the 1980s and 1990s. PNAS, 107(38): 16732–16737. 10.1073/pnas.0910275107

Gibson L, Lee T, Koh L. et al. 2011. Primary forests are irreplaceable for sustaining tropical biodiversity. Nature, 478: 378–381. 10.1038/nature10425

Hanski I. 2015. Habitat fragmentation and species richness. Journal of Biogeography, 42: 989–993. 10.1111/jbi.12478

Houghton R, Byers B, & Nassikas A. 2015. A role for tropical forests in stabilizing atmospheric CO2. Nature Climate Change, 5: 1022–1023. 10.1038/nclimate2869.

Lewis S, Edwards D, & Galbraith D. 2015. “Increasing human dominance of tropical forests.” Science, 349(625): 827–832. doi:10.1126/science.aaa9932

Liebsch D, Marques M, & Goldenberg R. 2008. How long does the Atlantic rain forest take to recover after disturbance? Changes in species composition and ecological futures during secondary succession. Biological Conservation, 141: 1717–1725

Myers S, & Patz J. 2009. Emerging threats to human health from global environmental change. Annual Review of Environment and Resources, 34: 223–252

Merritt M, Maldaner M, & de Almeida A. 2019. What Are Biodiversity Hotspots?. Frontiers of Young Minds, 7:29. doi:10.3389/frym.2019.00029

Mackey B, Kormos C, Keith H. et al. 2020. Understanding the importance of primary tropical forest protection as a mitigation strategy. Mitigation and Adaptation Strategies for Global Change, 25: 763–787. 10.1007/s11027-019-09891-4

Nunes C, Berenguer E, França F, Ferreira J, Lees A, Louzada J, Sayer E, Solar R, Smith C, Aragão L, Braga D, Camargo P, Cerri C, Oliveira R, Durigan M, Moura N, Oliveira V, Ribas, C, Vaz-de-Mello F, & Barlow, Jos. 2022. Linking land-use and land-cover transitions to their ecological impact in the Amazon. PNAS, 119. 10.1073/pnas.2202310119.

Pendrill F, Gardner T, Meyfroidt P, Persson U, Adams J, Azevedo T, Bastos-Lima M, Baumann M, Curtis P, De Sy V, Garrett R, Godar J, Goldman E, Hansen M, Heilmayr R, Herold M, Kuemmerle T, Lathuillière M, Ribeiro V, Tyukavina A, Weisse M, & West C. 2022. Disentangling the numbers behind agriculture-driven tropical deforestation. Science, 377

Pereira E, Ferreira P, Ribeiro L, Carvalho T, & Pereira H. 2019. Policy in Brazil (2016–2019) threaten conservation of the Amazon rainforest. Environmental Science & Policy, 100: 8–12. 10.1016/j.envsci.2019.06.001.

Projeto MapBiomas – Mapeamento Anual de Cobertura e Uso da Terra no Brasil-Coleção 9, acessado em 21.01.2024 através do link: https://brasil.mapbiomas.org/.

Renó V, Novo E, Suemitsu C, Rennó C, & Silva T. 2011. Assessment of deforestation in the Lower Amazon floodplain using historical Landsat MSS/TM imagery. Remote Sensing of Environment, 115: 3446–3456. 10.1016/j.rse.2011.08.008.

Ritchie H. 2023. Deforestation in the Amazon peaked decades ago. Can we get it to zero? Sustainability by numbers. https://www.sustainabilitybynumbers.com/p/amazon-zero-deforestation. Access it on. 21.01.2024

Silva J, Almeida R, & Carvalho L. (2023). An economic analysis of a zero-deforestation policy in the Brazilian Amazon. Ecological Economics, 203. 107613. 10.1016/j.ecolecon.2022.107613.

Silva C, Pessôa A, Carvalho N, et al. 2021. The Brazilian Amazon deforestation rate in 2020 is the greatest of the decade. Nature Ecology & Evolution, 5: 144–145. 10.1038/s41559-020-01368-x

Soterroni A, Mosnier A, Carvalho A, Câmara G, Obersteiner M, Andrade P, Souza R, Brock R, Pirker J, Kraxner F, Havlík P, Kapos V, zu Ermgassen E, Valin H, & Ramos F. 2018. Future environmental and agricultural impacts of Brazil’s Forest Code. Environmental Research Letters, 13. 10.1088/1748-9326/aaccbb

Souza C, Siqueira J, Sales M, Fonseca A, Ribeiro J, Numat I, Cochrane M, Barber C, Roberts D, & Barlow J. 2013. Ten-Year Landsat Classification of Deforestation and Forest Degradation in the Brazilian Amazon. Remote Sensing, 5(11): 5493–5513. 10.3390/rs5115493

Souza C, Oliveira L, Souza J, Ferreira B, Fonseca A, & Siqueira J. 2023. “Landsat Sub-Pixel Land Cover Dynamics in the Brazilian Amazon.”, Frontiers in Forests and Global Change 6:1294552. 10.3389/ffgc.2023.1294552.

Tonetti V, Niebuhr B, Ribeiro M, & Pizo M. 2022. Forest regeneration may reduce the negative impacts of climate change on the biodiversity of a tropical hotspot. Diversity and Distributions, 28: 2956–2971. 10.1111/ddi.13523

Watson J, Evans T, & Venter O. et al. 2018. The exceptional value of intact forest ecosystems. Nature Ecology & Evolution, 2: 599–610. 10.1038/s41559-018-0490-x

zu Ermgassen E, Adami M, Chaves M, Garcia A, & Hodel L. 2024. Halting the expansion of pasture in the Brazilian Amazon. One Earth, 7: 1923–1926. 10.1016/j.oneear.2024.10.012

